# Dynamics of supercoiled DNA with complex knots: large-scale rearrangements and persistent multi-strand interlocking

**DOI:** 10.1101/331314

**Authors:** Lucia Coronel, Antonio Suma, Cristian Micheletti

## Abstract

Knots and supercoiling are both introduced in bacterial plasmids by catalytic processes involving DNA strand passages. While the effects on plasmid organization has been extensively studied for knotting and supercoiling taken separately, much less is known about their concurrent action. Here, we use molecular dynamics simulations and oxDNA, an accurate mesoscopic DNA model, to study the kinetic and metric changes introduced by complex (five-crossing) knots and supercoiling in 2kbp-long DNA rings. We find several unexpected results. First, the conformational ensemble is dominated by two distinct states, differing in branchedness and knot size. Secondly, fluctuations between these states are as fast as the metric relaxation of unknotted rings. In spite of this, certain boundaries of knotted and plectonemically-wound regions can persist over much longer timescales. These pinned regions involve multiple strands that are interlocked by the cooperative action of topological and supercoiling constraints. Their long-lived character may be relevant for the simplifying action of topoisomerases.

## I. INTRODUCTION

The structural organization of bacterial plasmids is profoundly affected by homeostatic catalytic processes involving DNA such as transcription, replication and recombination. The best known example is the level of negative supercoiling, *∼* −5%, that is maintained by topoisomerases[1–3] and that creates an interplay between DNA twist (local winding around the double-helix axis) and writhe (winding of the double-helix axis around itself). Inter- and intra-molecular strand passages catalysed by topoisomerases can affect DNA entanglement too. The specific types of knots and links that can be introduced[4–8] or removed[8–10] by these mechanism, can vary with experimental conditions, the specific topoisomerase action on well-defined patterns of crossings [4] as well as DNA length[5–8, 10–12], which is a key determinant of knot complexity in viral [13–16] and eukaryotic DNA too[17].

The structural constraints associated to supercoiling and knotting have functional implications, too. Plectonemes, for instance, control the degree of branchedness of DNA rings [18–21] and this, in turn, affects the contact probability of *loci* at large genomic separations [18, 22–25]. In addition, the mechanics of superhelical stress, which can have long-range effects[26, 27], can facilitate the unzipping of AT-rich regions that are upstream of genes, and thus assisting the binding of promoters [26, 28–33].

Knots, on the other hand, have been mostly associated with detrimental functional effects, such as stalling DNA replication and transcription [5, 6, 34]. As a matter of fact, their statistically-inevitable emergence is counteracted by specific topoisomerases that can eventually remove them[10, 35, 36] via selective strand passages, arguably at hooked-juxtapositions that are characteristic of knots [37–40].

Intriguingly, recent modelling studies by Stasiak’s lab, have pointed to a primary role of supercoiling in the removal of DNA knots, too [41–43]. Stochastic simulations of coarse-grained DNA filaments tied in trefoil knots, the simplest non-trivial topology, have shown that the systematic accumulation of twist leads to a tightening, or localization, of the knot. This localization expectedly facilitates the local action (recognition and strand passage) of topoisomerases irrespective knot chirality [41, 42, 44].

These results add a novel appealing layer to the functional role of supercoiling. At the same time, its interplay with knots, especially the more complex ones reported in plasmids, is largely unexplored and key questions are still unanswered. For instance: how do knots more complex than trefoils affect the branchedness of supercoiled rings? Would the latter be increased by the large writhe of complex knots, or would it be suppressed by the topological constraints? Also, what is the effect of an intricate topology on the internal dynamics of plasmids, and how does it differ from the one of supercoiling? Do knots trap the system in long-lived states and, if so, what are their characteristics?

Here, we tackle these questions for 2kbp-long DNA rings using molecular dynamics simulations and oxDNA [45–47], an accurate model for DNA filaments based on mesoscopic representation of nucleotides and their interactions. This level of detail makes it possible to gather multi-ms trajectories for kbp-long DNAs while retaining the key structural details responsible for the frictional[48] and cholesteric effects[49] arising from self-contacts in the knotted or superhelical regions. Specifically, we focus on DNA rings with 5% negative supercoiling - typical of plasmids - and tied in 5-crossings knots (5_1_ and 5_2_ topologies), a complex form of entanglement previously reported in 4kbp-long pBR322 plasmids [5, 16].

We find several unexpected results. First, the conformational ensemble of knotted supercoiled rings is dominated by two states differing by knot length and number of plectonemes. Secondly, the spontaneous fluctuations between these states, which involve concerted changes of the knotted and plectonemically-wound regions, are fast, in that they occur on the same timescale of metric relaxation in unknotted rings, about 0.3ms. Strikingly, in spite of these large-scale variations, we observe that certain boundaries of the knotted region and of plectonemes persist throughout the 1.5ms-long simulated trajectories, and hence vary over much slower timescales. We show that these pinned boundaries involve multiple interlocked strands and that their slow evolution is exclusively due to the simultaneous action of knotting and supercoiling. We speculate that, in addition to previously established conformational features such as hooked juxtapositions or tight knots [38–40, 42], the long-lived character of these regions could aid the recognition, and hence knot simplication, by topoisomerases.

## II. MATERIALS AND METHODS

### A. Model

We considered 2kbp-long DNA rings with or without negative supercoiling and in three different topologies: 51 and 5_2_ left-handed knots as well as the unknot (0_1_ topology).

The rings were described with the oxDNA model [45–47], which provides an accurate mesoscopic description of double-stranded DNA. In the oxDNA model, each nucleotide is represented by three interaction centers (after the sugar, base and phosphate groups) with average (sequence-independent) parameters for base pairing, coaxial stacking, steric and electrostatic interactions. For the latter we used the default oxDNA paramerization for 1M monovalent salt.

### B. Initial setup

The initial conformations were generated with the following two-tier scheme. First we produced the centerline of the double-stranded DNA rings by using the Knot-Plot software (available at www.knotplot.com) to create smooth, symmetric forms of 0_1_, 5_1_ and 5_2_ knots, see Fig. 1A. To be consistent with experimental observations on pBR322 plasmids [5, 16] the chirality of the five-crossing knots was set to be left-handed, i.e. projected crossings have negative sign.

**FIG. 1.**
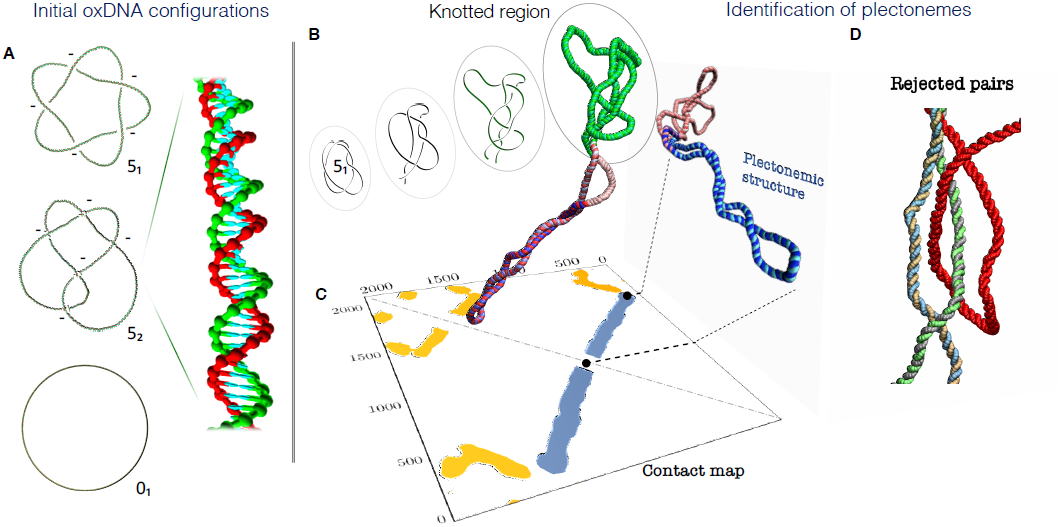
(a) Initial configurations of the supercoiled double-stranded DNA rings for the three considered topologies, 01, 51 and 52. The latter two are left-handed, i.e. the topological sign of their projected crossings is negative, as indicated. The 51 and 52 snapshots have been edited to highlight the over- and underpasses, see Supplementary Fig. S1 for the unedited versions. The mesoscopic structural representation of the oxDNA model is illustrated in the inset, which shows a magnified portion of one of the rings. The twist was uniformly adjusted for each of the three cases to yield the same level of negative supercoiling (-5%). (b-d) Identification of the knotted and the plectonemically-wound regions for a typical 51-knotted supercoiled conformation, shown in the foreground. (b) The knotted region (green) is the shortest portion that, after suitable bridging of the termini, has the same (51) topology of the entire ring. (c) The plectonemically-wound region (blue) is found by using the contact map to identify long superhelical regions ending in a short apical loop and that are free of *cis* or *trans* entanglement, as in the case of panel (d).

These circular centerlines, discretised in 2000 segments, were next turned into the oxDNA double-helical representation by a fine-graining procedure, where each segment was mapped into the six interaction centers of the two paired nucleotides [48], see inset in Fig. 1A. In this fine-graining procedure, the average twist between consecutive bases was adjusted differently for each topology to yield the sought level of supercoiling, as discussed in detail in ref. [42, 50]. Such twist adjustment is needed to account for the topological contribution to the writhe, i.e. the winding of the DNA centerline on itself. In fact, the canonical ensemble average of the writhe is usually different from zero for torsionally-relaxed knotted rings [51]. For each considered topological state we accordingly adjusted the twist uniformly to deal with the two different cases of torsionally-relaxed DNA rings and negatively supercoiled ones. In the latter case we set the relative amount of supercoiling equal to −0.05, the typical homeostatic level in bacterial plasmids.

### C. Molecular dynamics simulations

For each of the six combinations of knot types (0_1_, 5_1_ and 5_2_) and torsional states (relaxed and negatively-supercoiled) we collected ten different Langevin dynamics trajectories at *T* = 300K. The dynamical evolution was integrated with the LAMMPS package [52], using the implementation of Henrich *et al.* [53] and default values for the mass of the interaction centers, *m*, solvent viscosity, *η* and of the time step in the Langevin-<u>type</u> rigid-body integrator[53], 0.01*τ*_*LJ*_, where 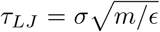 is the Lennard-Jones characteristic time and *ϵ* = *ι*_*B*_ *T* is the energy unit. Each trajectory had a typical duration of *∼* 2 × 10^7^ *τ*_*LJ*_. For the analysis, we omitted the initial relaxation phase of duration 10^6^*τ*_*LJ*_. The cumulative time span covered by all simulations was 1.2 × 10^9^*τ*_*LJ*_ and required about 1.4 × 10^6^ equivalent CPU hours on the Intel-based high-performance computing cluster (Ulysses) based in SISSA, Trieste.

In general, because of the concurrent presence of various dynamical regimes at different spatial scales, a correspondence between real time units and simulation time in coarse-grained models can be set only approximately. Here, we established the mapping *a posteriori* by matching the diffusion coefficient of the simulated DNA rings with the analogous experimental quantity. This conservative approach is expected to be more apt for the spontaneous dynamics of large systems than mappings based on the diffusivity of oligonucleotides[48]. Within the oxDNA setup, we found that the diffusion coefficients of supercoiled or torsionally-relaxed 2kbp-long unknotted rings at 1M monovalent salt are about *D*_theory_ *∼* 6.4 × 10^−4^*σ*^2^*/τ*_*LJ*_, where *σ* = 0.8518nm [45–47], see Supplementary Fig. S2. Experimental measurements for DNA rings of similar length yield *D*_exp_ *∼* 7 × 10^−12^ m^2^/s [54]. By equating *D*_theory_ and *D*_exp_, one therefore has *τ*_*LJ*_ *∼* 7 × 10^−11^s.

### D. Metric observables

As an overall metric observable we used the root-meansquare gyration radius,

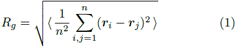

where *i* and *j* run over the *n* = 4000 nucleotides, ***r***_*i*_ is the position of the center of mass of the *i*th nucleotide, and the ⟨⟩ brackets denote the average over the configurations visited in the trajectories.

For the metric relaxation dynamics we computed the time-lagged autocorrelation function of *R*_*g*_

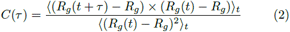

where *R*_*g*_(*t*) is the instantaneous gyration radius at time *t, τ* is the time lag, and ⟨⟩_*t*_ denotes the average taken over simulation time for the various trajectories. The characteristic timescale was computed as the integral of *C*(*τ*). To limit the effects of the noisy tail of *C*(*τ*) the integral was computed from *τ* = 0 up to when *C*(*τ*) drops below 10^−2^ for the first time.

To identify plectonemically-wound regions, if any, we generalized previous approaches[21, 55, 56] and used the multi-step strategy sketched in Fig. 1C-D and summarised below.

### E. Detection of plectonemes

We first constructed a contact map for the DNA cen-treline using a tolerant cutoff distance of 40*σ ∼* 32°Å, about three times larger than the typical superhelical diameter[12]. Next, to identify superhelical regions, we searched for clusters of contacts forming bands perpendicular to the contact map diagonal. These bands correspond to dsDNA stretches in spatial proximity and with opposite directionality (in an oriented ring).

For each band we identified its apex, which is the site, *i*, associated to the longest uninterrupted ladder of contacts {(*i*+Δ, *i −* Δ), (*i*+Δ+1, *i −* Δ *−* 1), (*i*+Δ+2, *i −* Δ *−* 2)}, *…* starting at a sequence separation, Δ, not larger than 400bp to avoid excessively large apical loops. At the same time trivial contacts due to short sequence separations Δ *<* 100bp are not counted. We note that at this stage, and at the next one too, there could be overlaps in the regions covered by different putative plectonemes. Within each band with Δ *<* 400bp, we then searched for the presence of tight contacts, a signature feature of superhelices. Specifically, we searched for the contacts at distance smaller than 7.5*σ*(*∼* 6Å), and took the contacting pairs with the smallest and largest sequence distance from the apex as the endpoints of the putative superhelical region. The putative plectoneme, formed by this region and the bridging apical loop was then checked to be free of entanglement by testing that progressively longer portions of the plectoneme had no physical linking [57] with the remainder of the ring. If this was not the case, the distal endpoints (those farthest from the apex) were progressively backtracked until the region became disen-tangled.

The plectoneme assignement was then carried out in an iterative non-overlapping manner, by ranking the putative plectonemes by the contour length of the superhelical region and disregarding instances where the latter was smaller than 300bp. The region with the longest super-helix was then assigned as the first plectoneme. Next, the distal endpoints of the remainder putative plectonemes (if any) were then backtracked to eliminate eventual overlaps with the sites assigned to the first plectoneme. The length ranking and selection was repeated and the second plectoneme was assigned so on, until exhaustion of the putative plectoneme set.

The iterative scheme allowed for the unsupervised detection of one or more plectonemes in practically all supercoiled configurations except for a small subset (0.5% of 5_1_-knotted instances and even smaller for 5_2_-knotted and unknotted ones) with non well formed superhelical regions or atypically large apical loops, see Supplementary Fig.S3.

### F. Topological observables

To locate the knotted region along the ring we used the bottom-up search scheme described in ref. [58]. The procedure consists of searching for the smallest portion of the ring (starting from portions of only few nucleotides and systematically expanding to longer ones) that, after closure, has the same topology of the entire ring. The topology was established using Alexander polynomials evaluated at *t* =*−* 1 and *t* = *−*2. For closing the considered portion we used the minimally interfering closure scheme[58], where the termini of the portion are bridged either with a straight segment or via a path involving the convex hull, depending on their proximity.

## III. RESULTS

### A. Conformational variability

We first discuss the conformational variability observed in simulations of 2kbp-long knotted DNA rings with the 5% negative supercoiling typical of bacterial plasmids. Respect to earlier studies on 3_1_-knotted DNAs, we stepped up the complexity and considered the 5_1_ and 5_2_ topologies, respectively a torus and twist knot. With these more complex knots we can explore the effects of a larger writhe and more numerous minimal crossings on the branchedness and dynamics of DNA rings. In addition, 5-crossings knots provide the simplest context for an equal footing comparison of torus and twist knots, which typically show different physical behaviour, from mechanical resistance to sliding friction to pore-translocation compliance [59, 60].

Typical snapshots, representative of the various degrees of branching found in the 5_1_- and 5_2_-knotted DNA rings, are given in Fig. 2A along with instances without knots (0_1_ case). The observed conformational variability is significant and is consistent with that recently reported for shorter unknotted plasmids based on cryo-em experiments and atomistic simulations [29]. It is interesting to examine the relationship between knottedness and the number of branches, or plectonemically-wound regions, because of the competing elements that govern it. The lobes, apices and clasps inherent to complex knots can favour plectonemes by serving as nucleation points, while the conformational restrictions of the topological constraints can inhibit plectonemes’ formation. The histograms in Fig. 2B clarify that, at this contour length, unknotted DNA rings are actually somewhat richer in plectonemes than knotted ones; in particular, instances with 3 or more plectonemes are practically found in unknotted rings only.

**FIG. 2.**
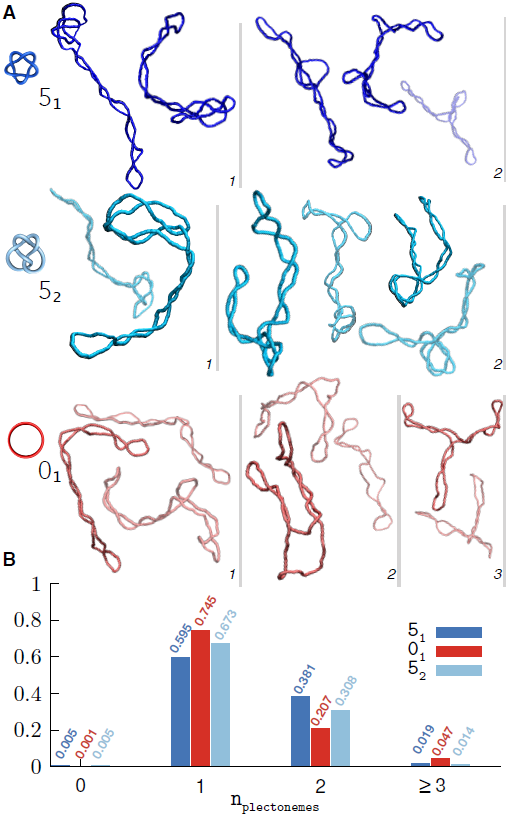
a) Typical snapshots of supercoiled DNA rings for the three considered topologies. The conformers are grouped by the number of plectonemes (in italics), which increases from left to right, and are shown in colours of different saturation for visual clarity. b) Normalised histogram of the number of plectonemes observed for each topology.

More conspicuous differences related to topology are found in the distributions of plectoneme lengths, *l*_*plc*_, and gyration radius, *R*_*g*_, see Fig. 3A,B. In particular, the conditional distributions of *l*_*plc*_ for the common single- and double-plectoneme states are little superposed for unknotted rings, but overlap substantially for knotted ones, see Supplementary Fig. S4 for the 5_2_ topology. The length of the plectonemes is also different across the 2kbp-long knotted and unknotted rings. For examples, plectonemes longer than 1500bp are common in uknotted rings but rare in 5_1_-knotted ones (50.7% and 0.05% of the populations, respectively). Conversely, conformers with only one plectoneme, and shorter than 1000bp, are uncommon in unknotted rings but abundant in 5_1_-knotted ones (6.3%and 22% of the populations, respectively).

**FIG. 3.**
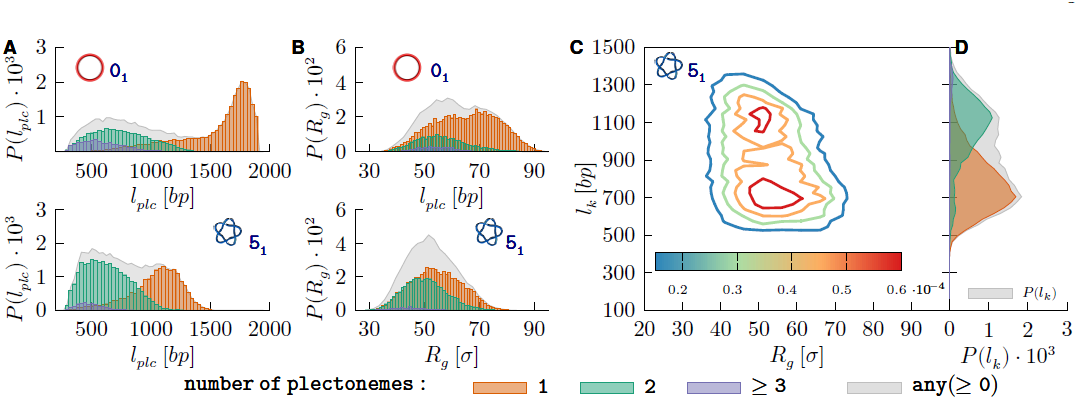
Probability distributions of the plectonemes’ length, *l*_*plc*_ and gyration radius, *R*_*g*_, for supercoiled rings with (A) unknotted and (B) 51 topologies. The conditional probabilities for 1 to 3 plectonemes are shown with coloured histograms, see legend, while the normalised combined distribution is shown in grey. (C) Normalised joint probability distribution of *Rg* and knot length, *l*_*k*_. (D) Marginal probability distribution of *l*_*k*_.

These differences could be of practical relevance, since they could be exploited in imaging, e.g. cryo-em, experiments to tell apart knotted from unknotted plasmids when supercoiling is present. Such discrimination is generally beyond the scope of gel electrophoresis, the method of choice for DNA topological profiling (but for a notable exception see [61]).

The interplay of knot length, *l*_*k*_, plectoneme length and gyration radius is presented in Fig. 3C-D. The joint probability distribution in panel C shows that *l*_*k*_ and *R*_*g*_ are anticorrelated, see also Supplementary Fig. S5, a property observed in other polymer systems too[62, 63]. What is specific of supercoiled knotted DNA rings is, instead, the presence of two peaks in the joint *l*_*k*_ *-R*_*g*_ distribution. The peaks’ origin is clarified by their marginal (projected) *l*_*k*_ distributions subdivided for number of plectonemes, see Fig. 3D. Specifically, the dominant peak, for *l*_*k*_ *∼* 700bp, is mostly associated to single-plectoneme states, while the peak at larger knot lengths (*l*_*k*_ *∼* 1100bp), corresponds to states with two or more plectonemes. Analogous results for the 5_2_ topology are shown in Supplementary Fig. S6.

The inverse correlation of *l*_*k*_ and the number of plectonemes is understood by noting that the knotted region, which is the shortest *uninterrupted* portion of the ring accommodating the essential crossings (the knotted core), must also include all intervening loops between the crossings except for the longest one. In supercoiled rings this remainder loop typically coincides with a plectoneme. Because the average plectoneme length decreases as their become more numerous, one has that *l*_*k*_ is shorter for states with a single plectoneme.

### B. Time evolution of metric and knot-related properties

The data shown in Fig. 4A are a kinetic counterpart to the static, or ensemble, view given above of the interplay of the knotted region, the number of plectonemes and the gyration radius.

**FIG. 4.**
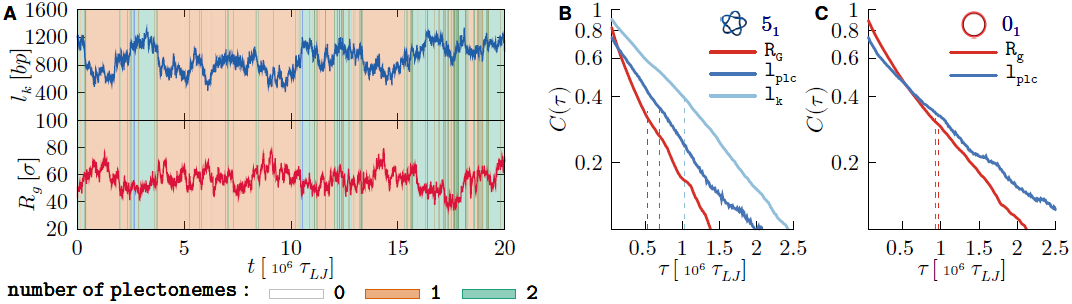
(A) Typical temporal traces of the length of the knotted region, *l*_*k*_ and the gyration radius, *R*_*g*_, from a trajectory of supercoiled 51-knotted ring. The background is coloured according to the instantaneous number of plectonemes, see legend. (B) Semi-log plot of the autocorrelation functions, based on data from all trajectories, of *R*_*g*_, *l*_*plc*_ and *l*_*k*_ of supercoiled 5_1_-knotted rings (B) and of *R*_*g*_, *l*_*plc*_ for unknotted ones (C).

One notes that over the typical duration of a trajectory (2.2 × 10^7^*τ*_*LJ*_ corresponding to about 1.5ms) both *l*_*k*_ and *R*_*g*_ have significant fluctuations, and clearly of opposite sign. These are accompanied by several changes in the number of plectonemes, as conveyed by the coloured background.

A more quantitative analysis of the characteristic timescales of these variations is given in Fig. 4B,C. These panels present the autocorrelation curves of *R*_*g*_, *l*_*k*_ and of *l*_*plc*_. The latter, was used in place of the number of plectonemes (on which it clearly depends) for its broader range of values, which makes it more amenable to the autocorrelation analysis. By integrating the autocorrelation curves, one has that the characteristic times of *l*_*k*_ and *R*_*g*_ are respectively equal to 1.03 × 10^6^*τ*_*LJ*_ and 0.53 × 10^6^_*τLJ*_, while for plectonemes’ length it is 0.69 × 10^6^*τ*_*LJ*_. Consistent with visual inspection, these timescales are all of the same order, about 10^6^*τ*_*LJ*_, which is much shorter than the duration of each simulated trajectory.

It is interesting that unknotted rings have a somewhat slower internal kinetics than knotted rings, cf. panels B and C in Fig. 4. In fact, the characteristic times of *R*_*g*_ and *l*_*plc*_ are, respectively 80% and 35% longer for unknotted rings (the same holds for *R*_*g*_ in the torsionally-relaxed case, see Supplementary Fig. S7). The result is not obvious, as one might expect a slower internal dynamics for knotted rings due to the friction of their self-contacts. It can be explained by considering that a finite portion of topological constraints necessarily use up a finite portion of the chain, and therefore knotted rings have a shorter effective contour length than unknotted ones and a smaller gyration radius too (see Fig. 3). This, in turn, reflects in a reduced breadth of the revelant con-formational space and hence a faster relaxation kinetics.

From the above analysis of overall metric and topological properties we conclude that, at physiological supercoiling, 2kbp-long knotted rings have enough conformational freedom to fluctuate spontaneously between two main states, related to the peaks in Fig. 3C, differing by knot size as well as the length and number of plectonemes. The characteristic timescale of these variations is *∼*3 × 10^6^*τ*_*LJ*_, corresponding to about 0.02ms, see Fig. 4A and Supplementary Fig. S8.

### C. Slowly-moving boundaries of the knotted region

To better understand the kinetics of the concerted variations of knot size and the number of plectonemes, we examined the time evolution of knotted and supercoiled regions, as in the kymographs of Fig. 5A, for other examples see Supplementary Fig. S9.

**FIG. 5.**
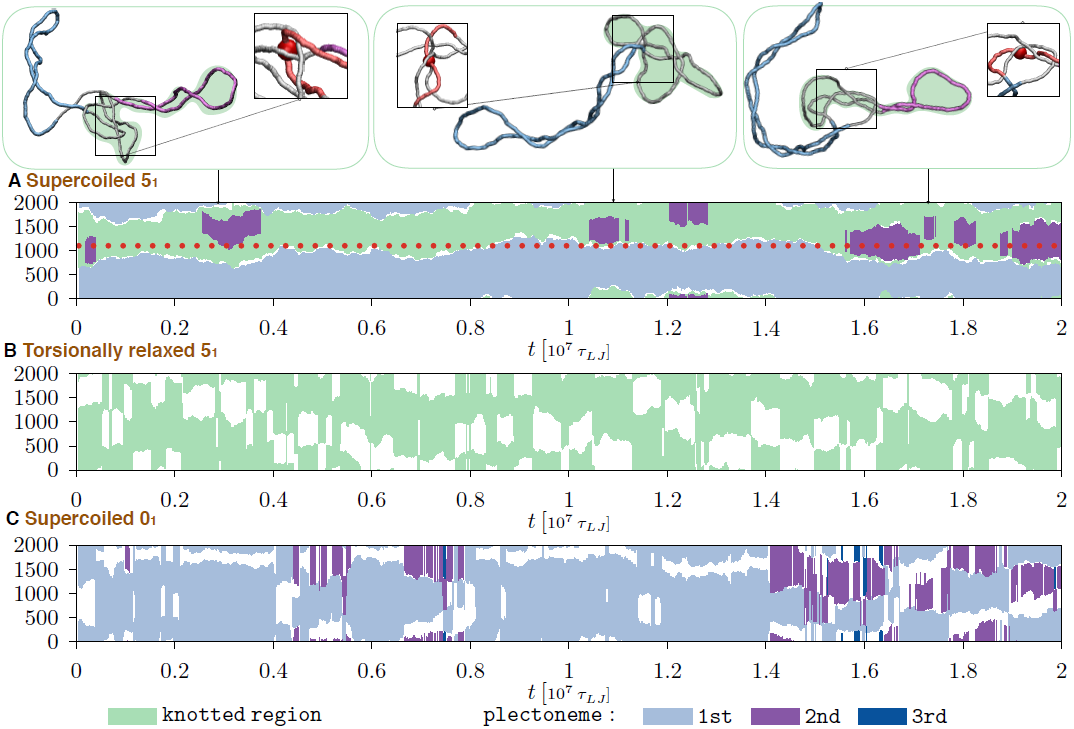
Kymographs showing the typical time evolution along the ring contour of knotted and plectonemically-wound regions, see legend for colour code. The three kymographs are for: (A) supercoiled 51-knotted rings, (B) torsionally-relaxed 51-knotted rings and (C) supercoiled unknotted rings. The boundaries of the knotted and the main plectonemically-wound regions of case are noticeable stabler than for case (B) and (C) due to persistent interlocking of multiple strands. This is illustrated in the snapshots above panel (A), where the same region at the knot-superhelix boundary (bp1000-bp1200, highlighted in red in the insets) remains entangled with other ring portions throughout the trajectory. The midpoint of this region is marked with a red bead in the insets and with a dotted red line in panel A.

The kymograph clearly indicates that there exists an additional relevant kinetic process besides those discussed before, namely a surprisingly slow stochastic motion of the knot along the ring contour.

Note, in fact, that the region covered by the knot at the beginning of the simulation in Fig. 5A (bp600-bp1800) still has not moved appreciably by the end of the trajectory (bp700-bp2000). Considering that the trajectory covers a timespans of 2 × 10^7^*τ*_*LJ*_ *∼* 1.5ms, it is clear that the contour motion of the knotted region occurs over timescales that are much longer than those discussed previously. These, we recall, were the relaxation time of the gyration radius and knot length, of about 10^6^*τ*_*LJ*_, and the changes between the tight and delocalised knotted states, which occur at intervals of about 3 × 10^6^*τ*_*LJ*_, see Fig. 4 and Supplementary Figs. S7 and S9.

We thus conclude that in supercoiled DNA rings, the contour motion of the knotted region is slower than these other processes by an order of magnitude (and likely more since the practical bound is the duration of the simulated trajectories).

Besides the knot, Fig. 5A shows the concurrent evolution of plectonemically-wound regions, too. Note that most of the time, there is a single long plectoneme that spans the ring portion complementary to the knotted region; but the latter can occasionally nest a plectoneme too. The kymograph clarifies that such nested instances have the following properties: (i) they occur in addition, and not in substitution, of the typically longer “dominant” plectoneme that complements the knotted region; (ii) their characteristic lifetime is 7 × 10^5^*τ*_*LJ*_ and (iii) consecutive appearances are separated by intervals of highly variable duration.

Overall, the several observed fluctuations of plectonemes’ number and length during the entire trajectory are in line with the relatively fast relaxation dynamics of *R*_*g*_ and *l*_*plc*_ of Fig. 4. Strikingly, these variations are accompanied by a noticeable persistence of the plectoneme boundaries, which mirrors the one of the knotted region. These features are ubiquitous across the collected trajectories for both 5_1_ and 5_2_ topologies, see Supplementary Fig. S9. One concludes that, irrespective of their torus (5_1_) or twist (5_2_) character, these supercoiled knotted rings have persistent boundaries between knotted and plectonemically-wound regions. This, in turn, poses the question of which of these two components, knots or plectonemes, is the primary cause for the slow evolution of these boundaries.

### D. Knotting and supercoiling are both required for persistent interlockings

To address this point, we decoupled entanglement and supercoiling by studying the time evolution of DNA rings where either the knot or supercoiling were present, but not both of them.

The results are shown in the kymographs of panels B-C of Fig. 5. Their properties are in stark contrast with those of panel A, where knotting and supercoiling were both present. In fact one observes that neither the boundaries of the knotted region (without supercoiling) nor those of plectonemes (without knotting) are inherently persistent. In fact, in both cases the boundaries vary on the same relatively fast timescales of the metric relaxation, i.e. *∼* 10^6^*τ*_*LJ*_.

We thus conclude that it is precisely the synergistic action of complex topology and supercoiling that is responsible for the locked boundaries, and the latter disappears when either of them is missing.

To clarify the mechanism underpinning this effect we inspected in detail the dynamical evolution of the rings. We thus established that the persistent boundaries correspond to specific points of tight and complex self-contacts of the knotted region.

A large number of self-contacting *loci* are clearly introduced by supercoiling in any DNA ring, regardless of its topological state. In knotted rings, due to the tightening of the intrinsic essential crossings, these *loci* typically involve several clasped or hooked double strands, as highlighted in the snapshots of Fig. 5A. We observed that, similarly to what happens during the pore translocation of knotted filaments [59, 60, 64, 65], the topological friction at these points is so high that the DNA strands are locally pinned while other parts of the chain can reconfigure.

This effect accounts for the observed separation of timescales between the metric relaxation time and the contour motion of the clasped points, at the boundary of the knotted region. Incidentally, we note that the relevance of these persistent regions of self-contacts reinforces *a posteriori* the necessity to use models, such as oxDNA, where the spatial description is sufficiently fine to capture the internal friction that develops when two tightly interacting DNA strands slide against each other. For this reason, we surmise that the observed sliding hindrance of contacting strands would be even higher in atomistic or more fine-grained models at this same high salt conditions. At the same time, the friction between DNA strands could be relieved by increasing their electrostatic repulsion with a lower salt concentration. We believe these would be worthwhile points to address in future modelling studies and possibly also experimentally, e.g. using setups akin to those of ref. [66].

Importantly, not all intrinsic (or essential) crossings of the knots create the persistent interlocking, but only a subset of them. This is visible in the snapshots of Fig. 5A where one notes that the two boundaries of the five-crossing knot are pinned by their high local physical entanglement and yet, the DNA strands can slide internally of the knotted region, despite the several points of pairwise contacts. It is precisely this internal sliding that creates the possibility for plectonemes to form transiently, but repeatedly, within the knotted region. As a matter of fact, the persistent interlocking appears in either of the two qualitatively different conformers populated by supercoiled knotted rings (i.e. those with local or non-local knots in Fig. 3.

In this regard, it is relevant to recall the seminal work of Liu *et al.* [38, 39], who pointed out that hooked DNA juxtapositions are an ideal target substrate of topoII enzymes because local strand passages at these point generally produces a simpler topology. The kinetic persistence of these multi-strand interlocking adds a novel temporal dimension to other, more thermodynamical effects of supercoiling, such as knot localization, that are credited to favour the local, yet globally-disentangling action, of topo II. The present results, in fact, complement the insight from earlier thermodynamic sampling[43, 44], by showing that once hooked or multiply-clasped juxtapositions are formed, they are long-lived. This, we speculate, is key for making such forms of local entanglement persistent enough to be recognsed and processed by topo II enzymes.

## IV. DISCUSSION

In summary, we used molecular dynamics simulation and the oxDNA mesoscopic model to study the effect of complex, five-crossing knots, on the conformational and kinetic properties of 2kbp-long plasmids with the typical 5% negative supercoiling found in bacterial plasmids. We particularly focussed on whether and how complex topologies, with their numerous points of high curvature and self-contact, can alter the branchedness of supercoiled plasmids and the dynamical evolution at long timescales.

On both accounts, we found that the interplay of knotting and supercoiling has major consequences that would have been difficult to anticipate *a priori*.

For the structural properties, we found that the conformational ensemble explored during the spontaneous dynamical evolution is largely dominated by two qualitatively-distinct states. They differ both by knot size and degree of branching. This fact, noteworthy *per se*, is accompanied by two intriguing kinetic effects. The first is that spontaneous fluctuations between these two states occur on timescales that are comparable to metric relaxation times of unknotted rings. The second is that certain boundaries separating the knotted and plectonemically-wound regions are very long-lived, and remain persistent over timescales that are much longer-arguably by at least an order of magnitude - than the metric relaxation times.

This complex phenomenology is shown to arise exclusively from the cooperative action of supercoiling and topological constraints; removing either of the two suffices to remove the persistent boundaries. The latter are shown to occur in correspondence of *loci* where multiple strands become interlocked. The interlockings have a local geometry that is analogous to the so-called hooked juxtapositions [38–40], argued to be ideal local targets for the topoisomerases’ knot simplifying action. We accordingly surmise that their long-lived character, besides their structural features, could also be instrumental to favour their recognition by topoisomerases. We believe this would be a noteworthy problem to address in future studies, for instance using mesoscopic models incorporating the interaction of DNA and proteins, which would be essential for a realistic description of DNA organization and processing *in vivo*. In addition, we expect that the incidence and long-lived character of multi-strand interlockings could also be probed with advanced single-molecule manipulation techniques such as pore translocation that, having been successfully applied DNA rings with either knots or supercoiling[48, 65, 67–71], is ideally suited to address their concurrent effects.

